# Quantitative proteomics indicate a strong correlation of mitotic phospho-/dephosphorylation with non-structured regions of substrates

**DOI:** 10.1101/636407

**Authors:** Hiroya Yamazaki, Hidetaka Kosako, Shige H. Yoshimura

## Abstract

Protein phosphorylation plays a critical role in the regulation and progression of mitosis. More than 10,000 phosphorylated residues and the associated kinases have been identified to date via proteomic analyses. Although some of these phosphosites are associated with regulation of either protein-protein interactions or the catalytic activity of the substrate protein, the roles of most mitotic phosphosites remain unclear. In this study, we examined structural properties of mitotic phosphosites and neighboring residues to understand the role of heavy phosphorylation in non-structured domains. Quantitative mass spectrometry analysis of mitosis-arrested and non-arrested HeLa cells revealed >4,100 and >2,200 residues either significantly phosphorylated or dephosphorylated, respectively, at mitotic entry. The calculated disorder scores of amino acid sequences of neighboring individual phosphosites revealed that >70% of dephosphorylated phosphosites exist in disordered regions, whereas 50% of phosphorylated sites exist in non-structured domains. A clear inverse correlation was observed between probability of phosphorylation in non-structured domain and increment of phosphorylation in mitosis. These results indicate that at entry to mitosis, a significant number of phosphate groups are removed from non-structured domains and transferred to more-structured domains. Gene ontology term analysis revealed that mitosis-related proteins are heavily phosphorylated, whereas RNA-related proteins are both dephosphorylated and phosphorylated, suggesting that heavy phosphorylation/dephosphorylation in non-structured domains of RNA-binding proteins plays a role in dynamic rearrangement of RNA-containing organelles, as well as other intracellular environments.

**Significance Statement:** Progression of mitosis is tightly regulated by protein phosphorylation/dephosphorylation. Although proteomic studies have identified tens of thousands of phosphosites in mitotic cells, the roles of them remain to be answered. We approached this question from the viewpoint of the higher-order structure of phosphosites. Quantitative proteomics and bioinformatic analyses revealed that more than 70% of mitotic dephosphorylation events occurred in non-structured regions. Non-structured regions of cellular proteins are attracting considerable attention in terms of their involvement in dynamic rearrangements of intracellular membrane-less organelles and protein assembly/disassembly processes. Our results suggest the possibility that a vast amount of mitosis-associated dephosphorylation/phosphorylation at non-structured regions plays a role in regulating the dynamic assembly/disassembly of intracellular architectures and organelles such as chromosomes and nucleolus.

## Introduction

Protein phosphorylation/dephosphorylation plays a critical role in a number of cellular processes, such as intracellular signaling, cell cycle regulation, and mitosis. Studies using mass spectrometry have identified tens of thousands of phosphorylation sites, resulting in the creation of an atlas of phosphorylation states and dynamic alterations in phosphorylation during the cell cycle (1), mitosis (2–4), and cell differentiation and development (3, 4).

A variety of structural biological approaches have been used to elucidate the effect of phosphorylation on the structure/function of substrate proteins. The addition of a phosphate group to the hydroxyl group of a target residue (Ser, Thr, or Tyr) can affect i) interactions with other proteins, ii) access of substrate molecules to the catalytic center of the enzyme, or iii) local internal energy, which can exert allosteric effects on other parts of the protein. Phosphorylation significantly affects stereo-specific interactions between the substrate and other molecules, thus ensuring tight regulation of biological reactions.

Recent bioinformatics studies revealed that phosphorylation occurs preferentially on residues in intrinsically disordered regions (IDRs) of proteins (5–7). Due to the absence of secondary and tertiary structures, IDRs are thought to function as flexible substrates for both phosphorylation and other post-translational modifications (6). Although the structural and functional significance of IDR phosphorylation remains to be fully elucidated, several possibilities have been proposed. For example, the addition of a phosphate group could reduce the flexibility of the IDR, thus facilitating specific interactions. Phosphorylation of a serine residue in the IDR of the transcription factor Ets1 alters the structure of the IDR and reduces the affinity of Ets1 for DNA (8). Phosphorylation of Ser19 in the IDR of the regulatory light chain of smooth muscle myosin induces the formation of an α-helix that activates an actin-dependent ATPase (9–11).

Another possible role for phosphorylation of IDRs could be to reduce the stereo-specificity between kinases and substrates. The human genome encodes ~520 different kinases and ~150 phosphatases, which phosphorylate and dephosphorylate, respectively, more than 40,000 sites in more than 10,000 proteins (12, 13). A single kinase (or phosphatase) can therefore phosphorylate (or dephosphorylate) many different substrates. Indeed, bioinformatic and biochemical studies have identified only weak consensus sequences for individual kinases (14) and almost no consensus around the target residues of phosphatases. The presence of a target residue in an IDR reduces the stereo-specificity of the enzyme-substrate interaction, thus enabling multiple enzyme-substrate combinations.

Mitosis is a critical cellular process tightly regulated by phosphorylation. Several thousand protein residues are known to be phosphorylated upon entry to mitosis (1, 2). A number of mitosis-associated kinases have been identified, including CDK1, Aurora kinase, Plk1, Bub1, and Haspin (reviewed in [17–21]). Dephosphorylation also plays an essential role during both mitotic entry and exit (22–25). More than 1,000 different phosphosites are known to be dephosphorylated during anaphase (22), and more than 500 are dephosphorylated upon entry to mitosis (2). For example, CDK1 is activated via dephosphorylation by Cdc25B/C (23, 24), and Cdc25C is partially activated by protein phosphatase 1 (23). These data indicate that a vast number of phosphate groups are transferred to and from substrate proteins in the early phase of mitosis.

Several questions regarding mitotic phosphorylation that have yet to be answered include why and how such a vast number of phosphate groups are transferred between different sets of proteins and whether these different mitosis-associated phosphosites differ structurally. Determining the distribution of phosphosites among structured and non-structured regions of substrate proteins is particularly important in terms of understanding the structural effects of phosphorylation/dephosphorylation. In this study, we therefore performed a phosphoproteomic analysis of cellular proteins to both quantify and compare individual phosphosites in mitosis and interphase. We also analyzed the structural properties (i.e., IDR or structured) of those sites. Our analysis of more than 6,000 phosphosites revealed a clear relationship between mitotic phosphorylation/dephosphorylation and IDRs.

## Results

### Quantitative proteomic analysis of phosphoproteins in mitosis

A comparative proteomics analysis was performed to examine phosphopeptides obtained from asynchronous and mitotic HeLa cells. Tryptic peptides from the two cell populations were labeled with tandem mass tag (TMT) reagents. LC-MS/MS analysis was performed after enrichment of phosphopeptides using TiO2. In total, 17,003 phosphopeptides were quantified (Fig. 1A), 15,368 of which were assigned to 3,701 proteins based on database searching. For quantitative analysis of phosphorylation, peptides with only a single phosphosite were extracted and assigned a mitotic abundance ratio, which provides an indication of enrichment in the specific phosphosite in mitosis relative to interphase (log_2_[M phase/asynchronous]). For phosphosites detected in more than two different peptides, the abundance ratio for each peptide was averaged. As a result, a total of 10,255 phosphorylation sites were quantified and subjected to further analyses, whereas 4,400 sites were detected only in peptides with multiple phosphosites. Of 10,255 phosphorylation sites, 9,656 were already registered in the PhophoSitePlus phosphoprotein database (25); thus, our dataset contained ≈600 previously unreported phosphosites.

**Figure 1.**
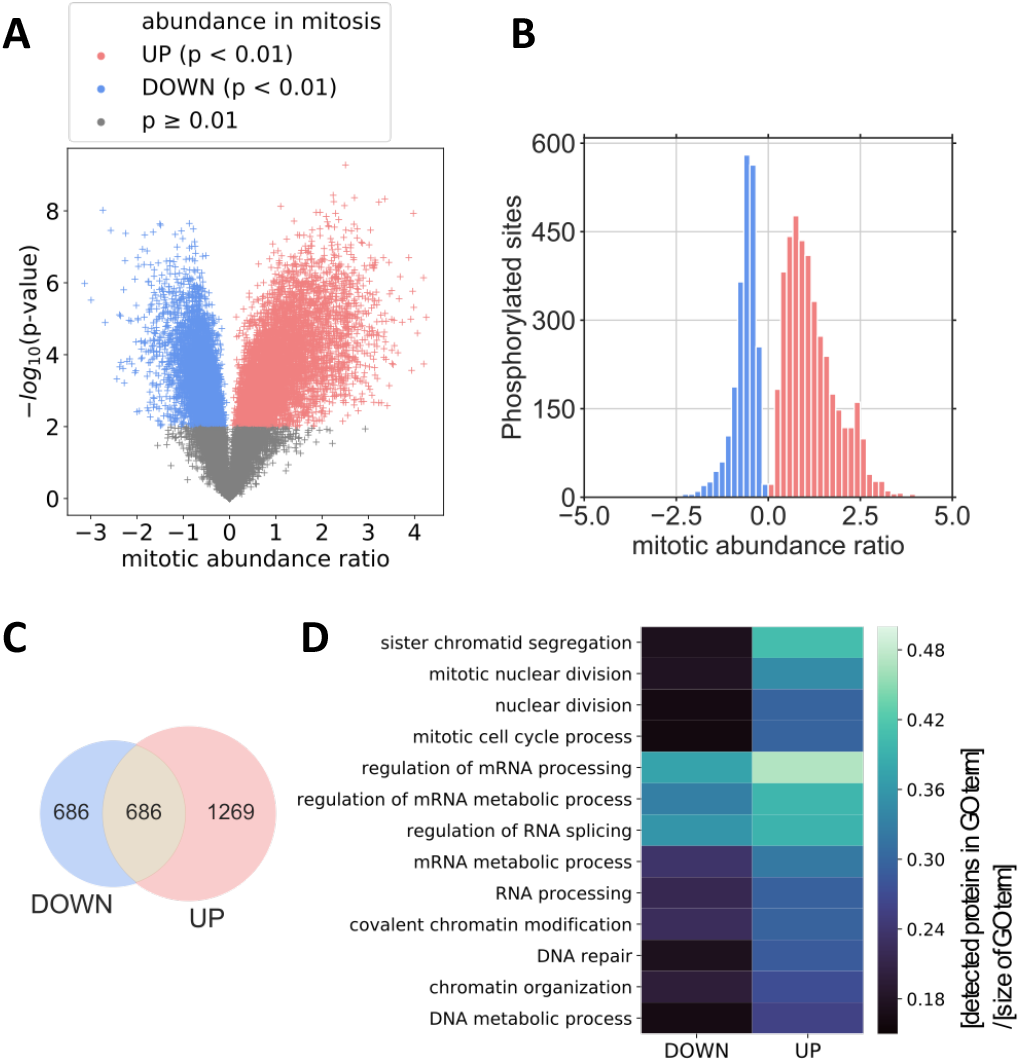
Quantification of mitotic phosphorylation by tandem mass tag analysis and gene ontology analysis on detected proteins. (A) Volcano plot of the TMT-analysis. The p-values and abundance ratios of individual phosphopeptides were plotted. Peptides with a −log_10_(p-value) <2 (grey) were eliminated from the subsequent analyses. Positive (red) and negative (blue) peptides were extracted and subjected to subsequent analyses. (B) Histogram of the abundance ratio. UP (red) and DOWN (blue) sites are distinguished. (C) Venn diagram of the number of proteins containing UP and/or DOWN sites. (D) Gene ontology analysis of phosphoproteins containing UP or DOWN sites. Biological process terms were obtained using DAVID. The number of proteins identified in this study was divided by the total number of proteins associated with each term. Shown are several terms relatively abundant for peptides with UP and DOWN sites.

To extract phosphosites specifically up- or down-regulated during mitosis, the p-values for individual peptides were plotted versus mitotic abundance. For a p-value threshold of 0.01 (Fig. 1A), two different phosphosite populations were extracted, one positive in terms of mitotic abundance (UP group) and the other negative (DOWN group) (Fig. 1B, Table S2). The UP group included 4,138 phosphosites distributed among 1,955 proteins, and the DOWN group included 2,249 phosphosites among 1,372 proteins. A significant number of proteins (686) contained both UP and DOWN sites (Fig. 1C). A small peak found at ≈2.4 (mitotic abundance ratio) (Fig. 1B) corresponded primarily to linker domains of C_2_H_2_ zinc finger proteins, which are known to be phosphorylated during mitosis (26). Several mitotic phosphosites known to be up-regulated during mitosis, such as S11 and S29 of histone H3.1 (27) and T161 of CDK1 (24), were identified in the UP group, with abundance ratios of 2.4, 1.3, and 0.8, respectively. Phosphosite T14 of CDK1, known to be down-regulated during mitosis (24), was detected in the DOWN group, with a mitotic abundance of −0.7.

Gene ontology (GO) analysis was performed for the UP and DOWN sites. Using DAVID, we obtained biological process terms (28, 29) and evaluated GO term enrichment in each group by dividing the number of proteins by the total number of proteins associated with specific GO terms (Fig. 1D). As expected, the UP sites were highly enriched in proteins related to mitotic chromosomes. Interestingly, both UP and DOWN sites were highly enriched in proteins related to RNA processing and splicing, suggesting that RNA-related proteins are regulated by both phosphorylation and dephosphorylation during mitosis (see *Discussion* section).

A larger number of UP sites than DOWN sites does not necessarily mean that there is a greater amount of phosphorylated proteins than dephosphorylated proteins in a mitotic cell. To examine this issue further, the phosphorylated proteins in cells in interphase and mitosis were quantified by dot-blot analysis using pIMAGO (30). Careful quantification revealed that 5.4 fmol of phosphate groups were attached to proteins in a single mitotic cell, versus 3.8 fmol in an asynchronous cell. These values correspond to 4.7 and 3.3 mM in the mitotic and asynchronous cells, respectively (the volume of a HeLa cell is reportedly 1.16 pL [35]) (Fig. S1A), indicating that mitotic cells contain 1.4 times more phosphate groups on proteins than asynchronous cells. These results were consistent with an observed decrease in the ATP concentration as determined using a FRET probe (Fig. S1B, C, D) (32, 33), and this decrease was inhibited by the universal kinase inhibitor staurosporine (Fig. S2A, B). The reduction in the ATP concentration was not due to cell swelling during mitosis (34, 35) (Fig. S1E), rearrangement of the actin cytoskeleton (Fig. S2C, D), or a decrease in the ATP supply (glycolysis and oxidative phosphorylation) (Fig. S2E-H). These results indicate that although both phosphorylation and dephosphorylation occur during mitosis, phosphorylation is the dominant process.

### Mitotic dephosphorylation occurs preferentially in non-structured regions

The relationship between mitosis-specific phosphorylation/dephosphorylation and the higher-order structure of polypeptides was also investigated. A previous study reported that phosphorylation of substrate proteins tends to occur in non-structured regions (IDRs) (5). We therefore evaluated the intrinsic disorder score of individual residues using the IUPred method (36, 37). In this study, the minimum length of the IDR was set to 30 amino acid residues (see *Materials and Methods*). Using this criterion, 23.4% of Ser/Thr residues among all cellular proteins are present in IDRs (Fig. 2A). In contrast, 59.6% of the 10,255 phosphosites quantified in our proteomic analysis were assigned in IDRs (Fig. 2A), demonstrating the strong likelihood of phosphorylation occurring in IDRs. As a comparison, other post-translational modifications were also subjected to the same analysis. As shown in Figure S3, the probabilities of ubiquitylation, methylation and acetylation sites existing in IDRs were lower than that of phosphorylation, confirming that this likelihood is specific to phosphorylation and not general tendency of post-translational modifications.

**Figure 2.**
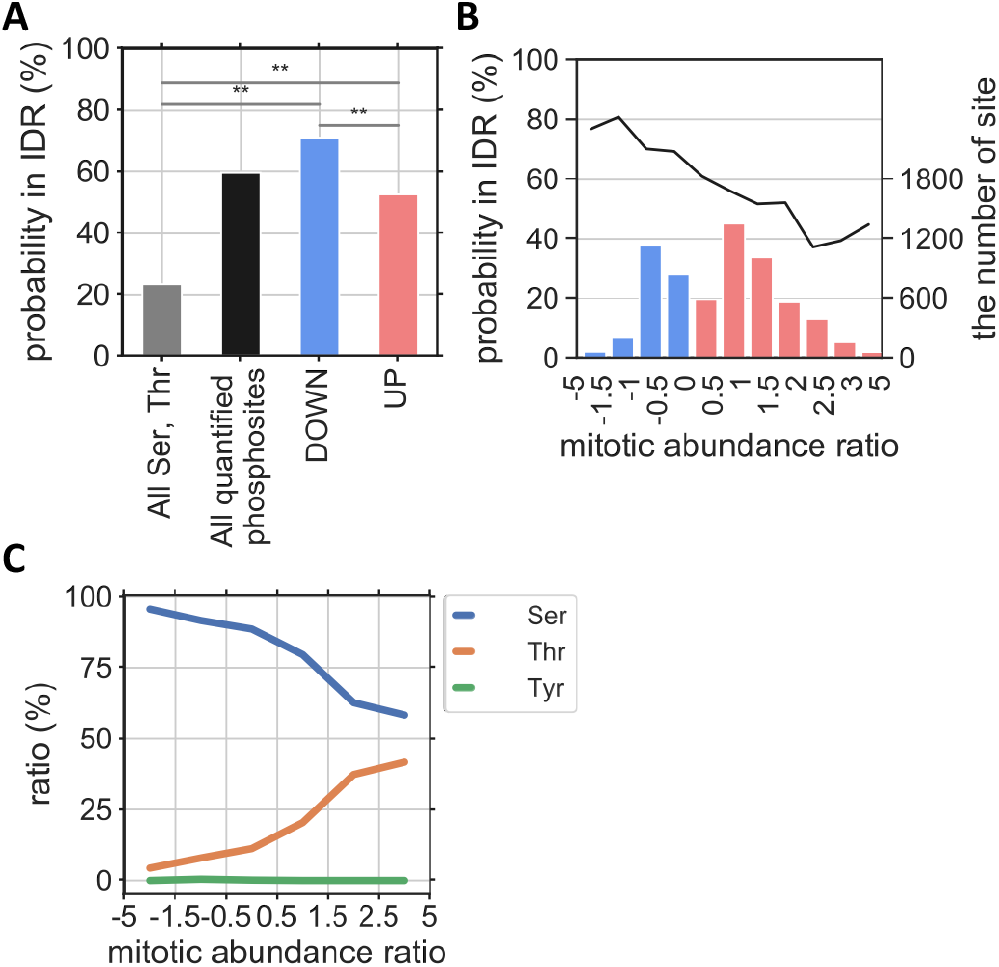
Structural properties and modified residues of mitotic phosphosites. (A) Probability of phosphosites existing in intrinsically disordered regions. The IDR probabilities of all Ser and Thr residues, phosphosites, UP sites, and DOWN sites are plotted. \*\**P* < 0.01. (B) Relationship between IDR probability and abundance ratio. (C) Relationship between phosphosite amino acid residue and abundance ratio.

We then examined the relationship between mitotic phosphorylation/dephosphorylation and IDRs. As shown in Figure 2A, 71.0% of DOWN and 52.6% of UP sites were found in IDRs. Interestingly, a strong inverse correlation between abundance ratio and IDR probability was found; the IDR probability gradually decreased from ~80 to 40% with increasing abundance ratio (Fig. 2B). These values were extremely high compared to the average for all Ser/Thr residues (23.4%). Collectively, these results demonstrate that phosphorylation generally occurs preferentially in IDRs, and this is particularly and significantly true for dephosphorylation upon entry to mitosis.

The specific phosphosite amino acid residue was also strongly correlated with the mitotic abundance ratio. As shown in Figure 2C, >90% of DOWN sites involved Ser, versus only ~7.8% for Thr residues. The percentage of Ser residues decreased as the mitotic abundance increased; 75.2% of UP sites were Ser, and 24.8% were Thr, indicating that mitotic dephosphorylation occurs primarily at Ser residues, whereas mitotic phosphorylation occurs at both at Ser and Thr residues. These results could be explained at least in part by the strong correlation between IDRs and DOWN sites, as Ser has a higher disorder probability than Thr. Alternatively, this could be associated with phosphatase preference. PP2A is known to prefer Thr over Ser and be inactivated at mitosis entry and re-activated in anaphase, as demonstrated previously (21, 22, 38, 39) (see *Discussion*).

### Mitotic phosphorylation–specific non-conventional sequence motifs

Next, we analyzed the sequence motifs specific to the UP and DOWN phosphosites. The UP and DOWN phosphosites were assigned as one of the following consensus sequences based on their flanking amino acids (from −6 to +6): “proline-directed” ([pS/pT]-P), phosphorylated by CDK and MAPK; “acidophilic” ([pS/pT]-X-X-[D/E], [pS/pT]-X-[D/E] or [D/E]-X-[pS/pT]), recognized by PLK1 and casein kinase; “basophilic” ([K/R]-X-X-[pS/pT] or [K/R]-X-[pS/pT]), phosphorylated by Aurora kinase, PKC, and PKA (2, 40); and “non-conventional” for those that did not match any of the three above categories.

As shown in Figure 3A and B, three conventional motifs (“proline-directed”, “acidophilic”, and “basophilic”) constituted 92.3% of DOWN sites, indicating that most mitotic dephosphorylation occurs at one of these conventional motifs. Interestingly, the amount of phosphorylation occurring at non-conventional motifs was higher for UP sites than DOWN sites (Figs. 3A and S4), suggesting the possibility of a mitosis-specific non-conventional motif. The conventional and non-conventional motifs also exhibited a clear contrast with regard to the relationship between the probability of phosphosites in IDR and mitotic abundance. As shown in Figure 3C, the probability of an IDR was inversely correlated with the mitotic abundance ratio for all three conventional motifs, as demonstrated in Figure 2A. In a clear contrast, the IDR probability in non-conventional motifs increased with increasing abundance ratio (Fig. 3C). The high IDR probability of UP sites was opposite to that of the conventional motifs. These results suggest that non-conventional motifs in IDRs are phosphorylated upon entry to mitosis.

**Figure 3.**
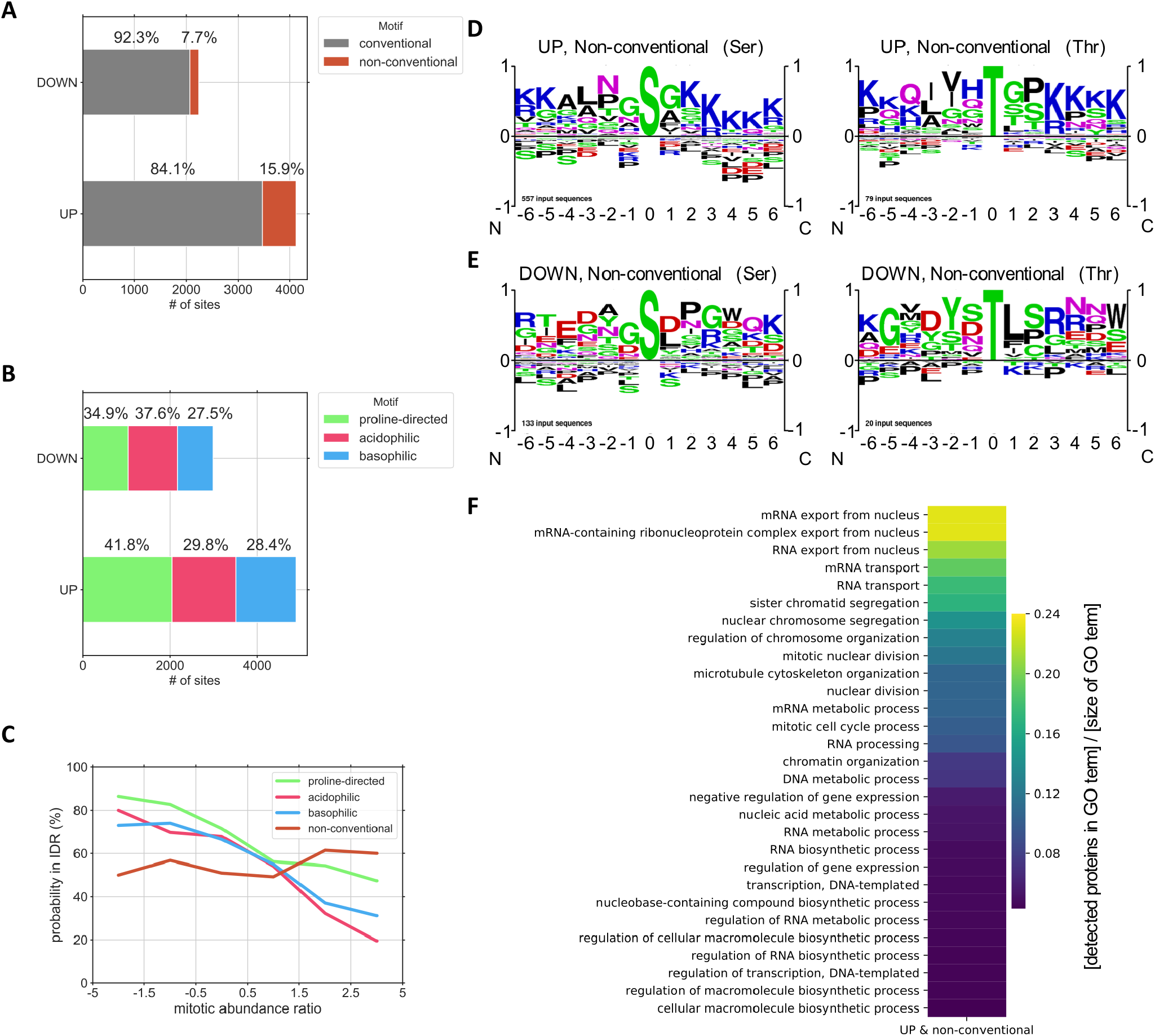
Neighboring sequences of mitotic phosphosites and the detected mitotic unconventional phosphorylation motif. (A, B) Ratio of conventional and non-conventional motifs (A) and ratio of individual conventional motifs (B) in UP and DOWN sites. (C) Relationship between IDR probability and abundance ratio for individual conventional and non-conventional motifs. (D, E) Logo analysis of residues flanking UP (D) and DOWN (E) sites in non-conventional motifs. Results for phospho-serine (left) and phospho-threonine (right) are shown. (F) Gene ontology (biological process) analysis of proteins with UP sites in non-conventional motifs. The number of proteins identified was divided by the total number of proteins associated with each term.

We then analyzed the amino acid sequences of “non-conventional” motifs. Logo analysis of 557 UP phospho-serine and 79 UP phospho-threonine sites in non-conventional motifs revealed a high frequency of basic residues (Lys or Arg) on the carboxylic side (+2 to +6) of the phosphosite (Fig. 3D). The presence of K/R at the +3 position was particularly notable. In contrast, no clear consensus was found regarding the amino-terminal side of the phosphosites, except for a weak consensus of a hydrophobic residue at the −2 position and K/R at the −6 position. This consensus was found to be distinct from any other known kinase motifs, including the conventional basophilic motif, which consists of a basic amino acid at the −2 or −3 position. In a clear contrast, no such consensus was found with respect to the DOWN sites; with no clear consensus for the 133 DOWN phospho-serine and 20 DOWN phospho-threonine sites in the non-conventional motifs (Fig. 3E). These results suggest that the non-conventional basophilic motifs (<u>N</u>on-conventional <u>b</u>asic residues at <u>c</u>arboxylic side motif (NBC motif)) are unique mitosis-specific phosphorylation sites.

GO term analyses were conducted for proteins carrying an UP site in the non-conventional motif (502 proteins). In contrast to the results of GO term analyses of all phosphoproteins with an UP site, proteins related to “mRNA export from the nucleus” were highly enriched in NBC motifs (24 of 502 proteins with non-conventional motifs [4.8%] vs. 50 of all 1,955 proteins with UP sites [2.6%]) (Fig. 3F). These proteins contain subunits of the nuclear pore complex (Nups), splicing factors, and other RNA-binding proteins. Nups and proteins with transcription regulator activity are known to carry a substantial number of IDRs (41, 42). These results, together with those regarding dephosphorylation of RNA-related proteins (Fig. 1D), suggest that phosphorylation/dephosphorylation of RNA-related proteins play important roles in the progression of mitosis.

## Discussion

In this study, we conducted a comparative proteomic analysis of amino acid residues phosphorylated in cells undergoing mitosis and interphase cells using a TMT-6plex labeling technique that provides a ratio of the amounts of individual peptides in six samples with high precision (43). Using this approach, we extracted two different populations of phosphopeptides, one up-regulated and the other down-regulated upon entry to mitosis. Bioinformatic analyses of these phosphopeptides revealed a clear correlation between phosphorylation/dephosphorylation and the IDR probability of the substrate. The most striking outcome of our analysis was that although phosphorylation generally tends to occur in IDRs, this trend is more common with mitotic dephosphorylation, as >70% of DOWN sites were found in IDRs, versus only 50% of UP sites (Fig. 2A, B). These data indicate that a significant number of phosphate groups are removed from IDRs and introduced into more structured regions upon entry to mitosis. Although we did not elucidate the significance of this translocation of phosphate groups from IDRs to structured regions in this study, several intriguing clues can be discerned from analyses of flanking amino acid sequences and GO term analysis: i) most mitotic dephosphorylation occurs at Ser residues (91.9 %), whereas phosphorylation occurs at both Ser and Thr residues (75.2%, 24.8%, respectively) (Fig. 2C); ii) proteins involved in mitotic chromosome segregation are heavily phosphorylated, whereas proteins related to RNA splicing and metabolism are both dephosphorylated and phosphorylated (Fig. 1D); and iii) non-conventional basic motifs are preferentially phosphorylated during mitosis (Fig. 3A, D, E). These intriguing results suggest a correlation between mitotic phosphorylation and dynamic higher-order assembly/disassembly of biomolecules (proteins, DNA, and RNA). Below, we discuss the potential significance of such dynamic behavior of biomolecules in the context of cellular function.

One effect of phosphorylation in IDRs could be to facilitate the transition of higher-order structures (8, 44, 45). Phosphorylation in IDRs is known to cause disordered-to-ordered and ordered-to-disordered transitions. For example, phosphorylation of 4E-BP2 induces the formation of a four-stranded β-sheet, thus preventing the binding of eIF4E (46). Phosphorylation of phospholamban at Ser16, in contrast, disrupts the higher-order structure, in turn reducing the inhibitory effect of phospholamban on sarco(endo)plasmic Ca-ATPase activity (47, 48).

Another possible role for phosphorylation in IDRs is regulation of the phase transition of biomolecules. Non-structured polypeptides of IDRs have been demonstrated to play key roles in liquid-liquid phase separation (49). It is driven by a number of promiscuous interactions to assemble a number of molecules. Other studies demonstrated that phase transition of protein liquid droplets is induced by phosphorylation/dephosphorylation (50–52). For example, liquid-liquid phase separation of a complex of a positively charged artificial peptide and RNA is promoted by dephosphorylation of the peptide (53). A study using a short hydrophobic peptide that forms a hydrogel demonstrated that reversible gel-sol transition is induced by cyclic phosphorylation-dephosphorylation by kinases and phosphatases (54). The peptide hydrogel is transformed into solution upon addition of kinase and ATP to the gel and re-solidifies when phosphatase is added to the solution. Such phenomena have also been observed in living cells. The assembly/disassembly of RNA granules is regulated by phosphorylation/dephosphorylation of one of the subunits (55). Although the mechanism is not fully understood, available evidence suggests a possible effect of phosphorylation/dephosphorylation on the dynamic assembly/disassembly transition of biomolecules upon entry to mitosis. It is possible that phosphate groups in IDRs function in weakly assembling the proteins during interphase, and their removal (dephosphorylation) could induce final assembly of the proteins into a stable complex. Alternatively, mitotic phosphorylation could solubilize/disassemble the protein complex, and dephosphorylation during anaphase could reassemble the complex. Indeed, phosphorylation of proteins in splicing speckles, pericentriolar-satellites, and stress granule by DYRK3 results in disassembly of these membrane-less organelles during mitosis (56). Such phosphorylation-dependent regulation of protein assembly/disassembly depends on the sequence of amino acid residues flanking the substrate residue. A previous report demonstrated that the charge pattern—rather than the net charge—is most important for phase separation (57). The addition of a negative charge due to phosphorylation could enhance or disrupt the charge pattern and thus affect the progress of phase separation.

Our GO term analysis of UP and DOWN phosphosites revealed that RNA-related proteins are preferentially dephosphorylated and phosphorylated during mitosis (Fig. 1D). This result is intriguing in the context of phosphorylation-dependent regulation of phase transition in intracellular compartments. A recent study demonstrated that the structure and function of several intracellular compartments are maintained by phase separation, in which RNA plays a role in the molecular assembly (58). P-granules are cytoplasmic compartments in the germ line of *C. elegans*, and they are considered liquid-like condensates containing mRNA and RNA-binding proteins that function in posttranscriptional regulation (59, 60). Protein components of P granules, such as PGL-3 and LAF-1, form liquid droplets *in vitro*, and mRNA affects the dynamics of these proteins within the droplets (64, 65). In the nucleolus, which is also considered a compartment, proteins and RNAs assemble via a mechanism similar to phase separation (63). The nucleolar protein nucleophosmin forms droplets with rRNA *in vitro* (64). Phosphorylation of Thr^199, 219, 234, 237^ by CDK1 reduces the affinity for rRNA (65), and replacement of these Ser residues with Glu partially abolishes nucleolar localization in HeLa cells (66), suggesting that phosphorylation of nucleophosmin controls the dynamics of proteins and RNAs within the nucleolus. It is possible that the removal/addition of a large number of phosphate groups from/to IDRs of RNA-related proteins induces dynamic rearrangements of the nucleolus and other RNA-containing nuclear speckles upon entry to mitosis, resulting in disruption of these organelles and the release of a large number of RNA molecules into the cytoplasm. This would dramatically alter the intracellular environment and could affect the assembly/disassembly dynamics of other protein complexes, such as chromosomes, which are known to contain pre-ribosomal RNA and nucleolar proteins on the surface (67, 68). It is also possible that chromosome condensation could be induced by the orchestrated effects of phosphorylation of chromosome-related proteins (Fig. 1D) and dramatic RNA-induced changes in the intracellular environment. Further study will be required to resolve this issue.

An unexpected result of the present study was that most dephosphorylation upon entry to mitosis was found to occur at Ser residues, whereas phosphorylation was found to occur at both Ser and Thr residues (Fig. 2C), suggesting that Thr phosphorylation is a mitosis-specific event. It is possible that such Thr phosphorylation of substrate proteins is coupled with dephosphorylation during anaphase and therefore temporary during early mitosis. There are several lines of experimental evidence that support this possibility: i) dephosphorylation during anaphase and telophase occur preferentially at Thr residues in HeLa cells (22, 38); ii) in budding yeast, Thr residues are heavily phosphorylated upon entry to mitosis and dephosphorylated during anaphase by PP2A^Cdc55^, an orthologue of PP2A^B55^ (21, 39, 69); iii) biochemical analyses revealed that PP2A^B55^, a major anaphase-associated phosphatase, preferentially dephosphorylates Thr residues over Ser residues (70); and iv) a comparison of our dataset of UP sites with a previously reported dephosphorylation dataset (22) revealed that 29.9% of Thr and 9.4% of Ser residues detected in the both datasets are dephosphorylated during anaphase. These data suggest that Thr phosphorylation during mitosis is temporary and functions to regulate reactions that are tightly tuned temporally during mitosis. The combined activities of kinases and phosphatases enable such tight regulation. Although the structural background of Thr-specific reactions has yet to be elucidated, it must be involved in early mitotic events such as chromosome condensation.

We identified mitosis-specific phosphorylation motifs similar to conventional basophilic motifs but different in the position of basic residues (Fig. 3D). Although distinct from any known phosphorylation motifs, several candidates were found in a phosphoproteomic database (2). PKC is known to function in mitosis (71, 72) and phosphorylate Ser residues near basic amino acids. However, the consensus sequences of several PKC subtypes contain basic residues on the C-terminal side as well (40). PKCε plays a role in resolving mitotic DNA catenation (73). Our GO term analysis revealed that proteins related to sister chromatid segregation are enriched in proteins carrying NBC motifs. Another potential candidate is AMPK, which is activated when AMP levels increase and ATP levels decrease (74) and known to phosphorylate mitosis-related proteins (75). Although the primary consensus motif for AMPK is basophilic, a peptide with K/R at the +3 position, which is the same as the NBC motif, is also phosphorylated *in vitro* (76). Of 243 proteins identified as substrates of AMPK to date (75), 17.3% (42 proteins) were found to contain UP sites in the NBC motif in our study (Table S2). Furthermore, it is intriguing that the intracellular ATP level decreased during early mitosis (Fig. S1B-D). These results suggest that mitosis is regulated by an ATP-dependent regulatory mechanism; decreasing intracellular ATP levels resulting from the activity of conventional kinases upon entry to mitosis may activate AMPK, which then phosphorylates different sets of substrate proteins to induce the progression of mitosis. The activity of AMPK increases during early mitosis and then decreases when cytokinesis begins (75), in good agreement with observed ATP levels during mitosis (Fig. S1C).

## Materials and Methods

### Cell culture and synchronization

HeLa cells were cultured in Dulbecco’s modified eagle medium (DMEM) (Sigma-Aldrich) with 10% fetal bovine serum (FBS) (GIBCO) at 37°C and 5% CO_2_. For mitosis-arrested cells, cells were treated first with 2 mM thymidine (Sigma-Aldrich) for 18 h, washed with PBS, and released into DMEM with 10% FBS for 1 h. Following treatment with 0.2 μM nocodazole for 10 h, 80% synchronization of HeLa cells in mitosis was achieved.

### TMT labeling and mass spectrometry

Asynchronous and mitosis-arrested HeLa cells in 100-mm cell culture dishes (Corning) were washed twice with 5 mL of ice-cold Hepes-saline (20 mM Hepes-NaOH [pH 7.5], 137 mM NaCl). Next, 0.75 mL of guanidine hydrochloride buffer (6 M guanidine hydrochloride, 100 mM Tris-HCl [pH 8.0], 2 mM DTT) was added, and cell lysates were prepared in triplicate and frozen in liquid nitrogen. The lysates were dissolved by heating and sonication, followed by centrifugation at 20,000 × *g* for 15 min at 4°C. The supernatants were reduced in 5 mM DTT at room temperature for 30 min and alkylated in 27.5 mM iodoacetamide at room temperature for 30 min in the dark. Proteins were purified by methanol/chloroform precipitation and solubilized by addition of 25 μL of 0.1% RapiGest SF (Waters) in 50 mM triethylammonium bicarbonate. After repeated sonication and vortexing, the proteins were digested with 2 μg of trypsin/Lys-C mix (Promega) for 16 h at 37°C. The peptide concentration was determined using a Pierce quantitative colorimetric peptide assay (Thermo Fisher Scientific). Approximately 150 μg of peptides for each sample was labeled with 200 μg of TMT-6plex reagent (Thermo Fisher Scientific) for 1 h at room temperature. After the reaction was quenched with hydroxylamine, all TMT-labeled samples were pooled and acidified with trifluoroacetic acid (TFA). Phosphopeptides were enriched using a 50-mg column of Titansphere Phos-TiO (GL Sciences) in accordance with the manufacturer’s instructions and then fractionated using a Pierce high-pH reversed-phase peptide fractionation kit (Thermo Fisher Scientific). Eight fractions were collected: 5, 7.5, 10, 12.5, 15, 17.5, 20, and 50% acetonitrile. Each fraction was evaporated in a SpeedVac concentrator and dissolved in 0.1% TFA.

### Data analysis

Data analysis was performed using Python2 or 3 and the accompanying libraries (Numpy, Scipy, Pandas, Matplotlib, Seaborn) using Jupyter (IPython) Notebook. The latest version of the phosphosites dataset from PhosphoSitePlus (25) (Wed Nov 08 15:57:30 EST 2017) was used. For GO term analysis, DAVID 6.8 (28, 29) was used. The term “BP_5” was obtained as a biological process term applying *Homo sapiens* as the background. The number of proteins in our dataset was divided by the total number of proteins for each GO term to evaluate the abundance of the term. For IDR analysis, the IUPred method (36, 37) was used. Position-specific estimations of energies of each residue were calculated based on the method described by Dosztányi using Perl script. Residues with an energy greater than −0.203 [aeu] were defined as intrinsically disordered residues according to the IUPred method. Contiguous amino acid sequences of more than 30 intrinsically disordered residues were regarded as intrinsically disordered regions. LOGO analysis of the neighboring amino acids of phosphosites was performed using the “PSP production” algorithm in PhosphoSitePlus (25).

## Acknowledgments

This study was supported financially by The Sumitomo Foundation Grant for Basic Science Research Projects (Grant Number 150852) for S.H.Y and JSPS Grant-in-Aid for JSPS Fellows (Grant Number 17J09002) for H.Y. We thank T. Oda for the assistance in the analysis of IDR.

## Supplemental Figure, Table and Legends

**Figure S1.**
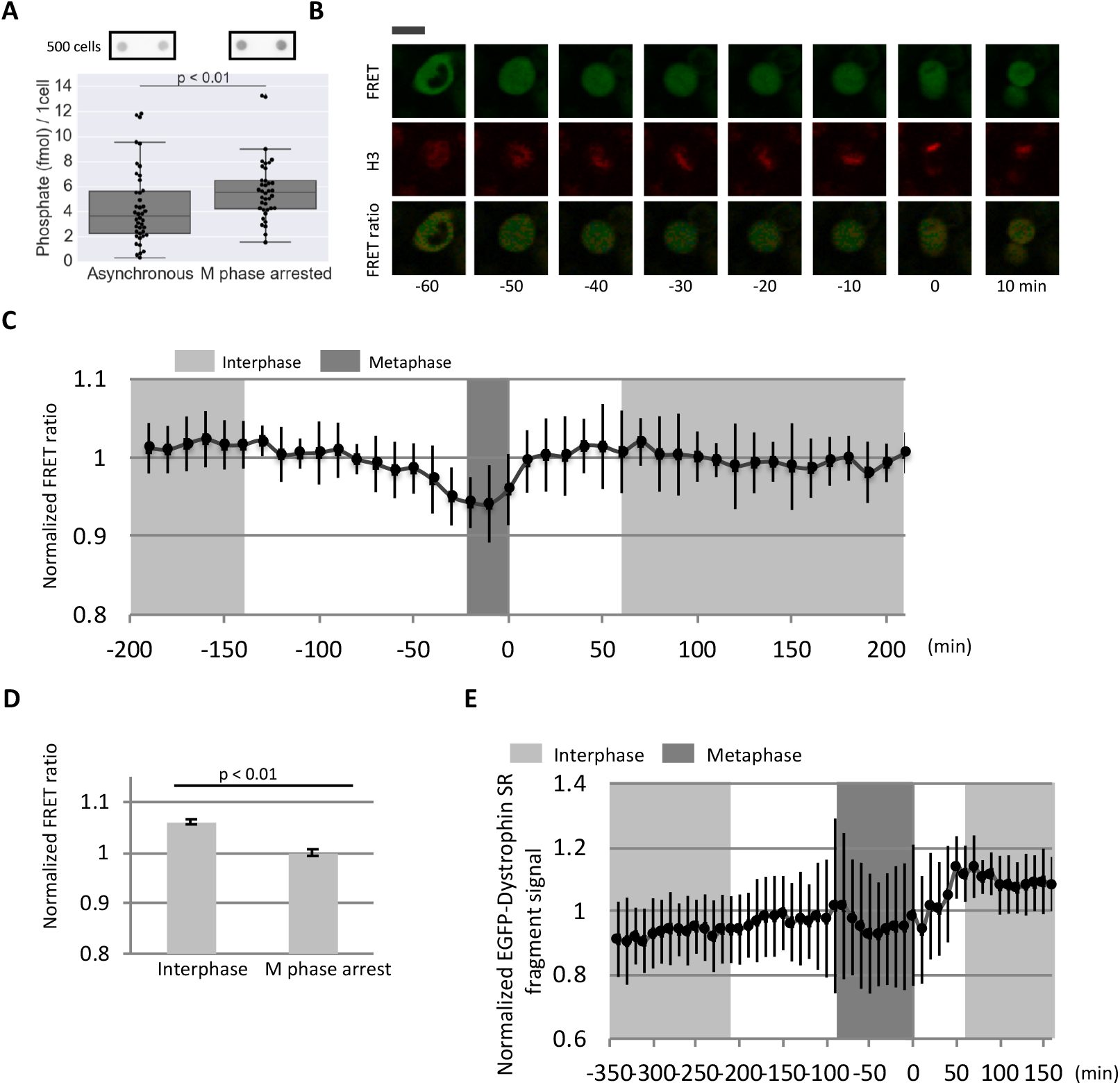
Total phosphoprotein amount increases and intracellular ATP level decreases during mitosis. (A) The amount of phosphate on proteins in asynchronous and nocodazole-treated HeLa cells was quantified using pIMAGO and streptavidin HRP conjugate. Dot-blot analysis of the lysate of ~500 cells is shown. The data points were obtained from three independent experiments, and outliers were eliminated in each experiment based on the inter-quartile range. Significance was assessed using the Mann–Whitney *U*-test. (B, C) Quantification of the cytoplasmic ATP level during mitosis. HeLa cells expressing ATeam and mPlum-histone H3 were observed by live-cell time-lapse imaging. Representative fluorescence images of FRET signal, mPlum signal, and acceptor/donner ratio are shown. Scale bar: 20 μm (B). Time 0 was defined as the time chromosomes began to segregate at anaphase onset. The acceptor/donner ratio was quantified in images, normalized to that of cells in interphase, and plotted versus time (C). Data were collected from 16 cells, and each data points corresponds to at least 8 cells. Error bars represents S.D. (D) Comparison of cytoplasmic ATP level between non-treated and nocodazole-treated HeLa cells. HeLa cells expressing ATeam were treated with 2 mM thymidine for 18 h and released to DMEM with 10% FBS for 1 h. The cells were then treated with or without 0.2 μM nocodazole for 10 h and observed by confocal fluorescence microscopy. The acceptor/donner ratio was quantified in the obtained images and summarized. Data were collected from more than 100 cells. Values were normalized to those of nocodazole-treated cells. A reduction in signal in mitosis was observed, as was the case in living cells shown in B and C. Error bars represent 95% CI. Significance was assessed using Welch’s *t*-test. (E) The experiment described in Figure S1B and C was performed with EGFP-tagged dystrophin. Data were collected from 13 cells, and each data point corresponds to at least 7 cells. Error bars represent S.D.

**Figure S2.**
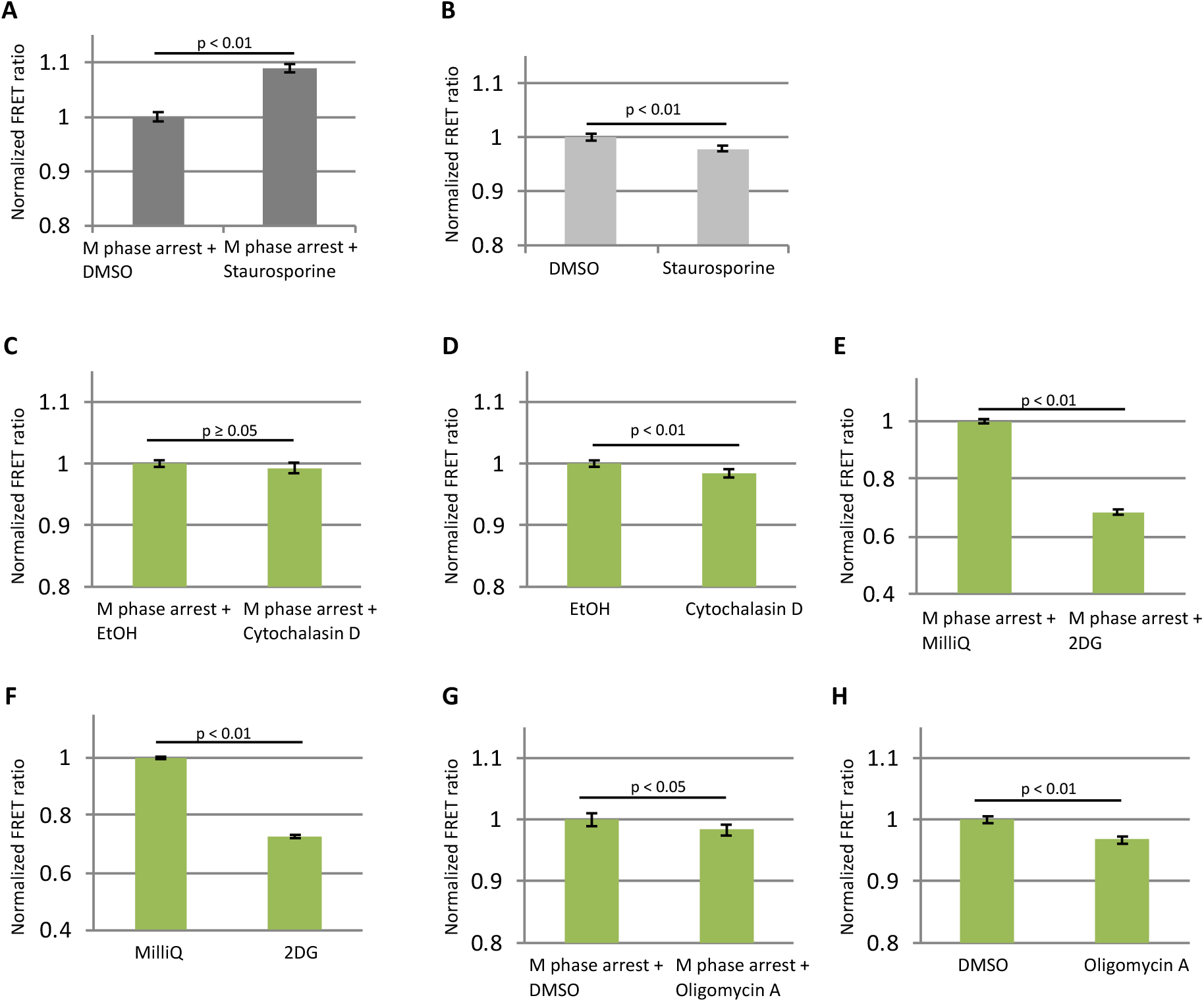
Protein phosphorylation decreases the intracellular ATP level during mitosis. The experiment described in Figure S1D was performed in the presence of various inhibitors in M-phase–arrested cells (A, C, E, G) and non-synchronized cells (B, D, F, H). For M-phase arrest, HeLa cells were treated with thymidine for 18 h, and after incubation with normal medium for 1 h, the cells were treated with 0.2 μM nocodazole for 10 h. (A, B) Cells were treated with 1 μM staurosporine, a universal inhibitor of kinases for 1 h. (C, D) Cell were treated with 1 μM cytochalasin D, an inhibitor of actin polymerization, for 1 h. (E, F) Cells were treated with 10 mM 2-deoxy-D-glucose (2DG), an inhibitor of glycolysis, for 1 h. (G, H) Cells were treated with 12.6 μM oligomycin A, an inhibitor of oxidative phosphorylation, for 1 h. The acceptor/donner ratio was quantified in the obtained images and normalized to that in the absence of the inhibitor. Error bars represent 95% CI. Significance was assessed using Welch’s *t*-test.

**Figure S3.**
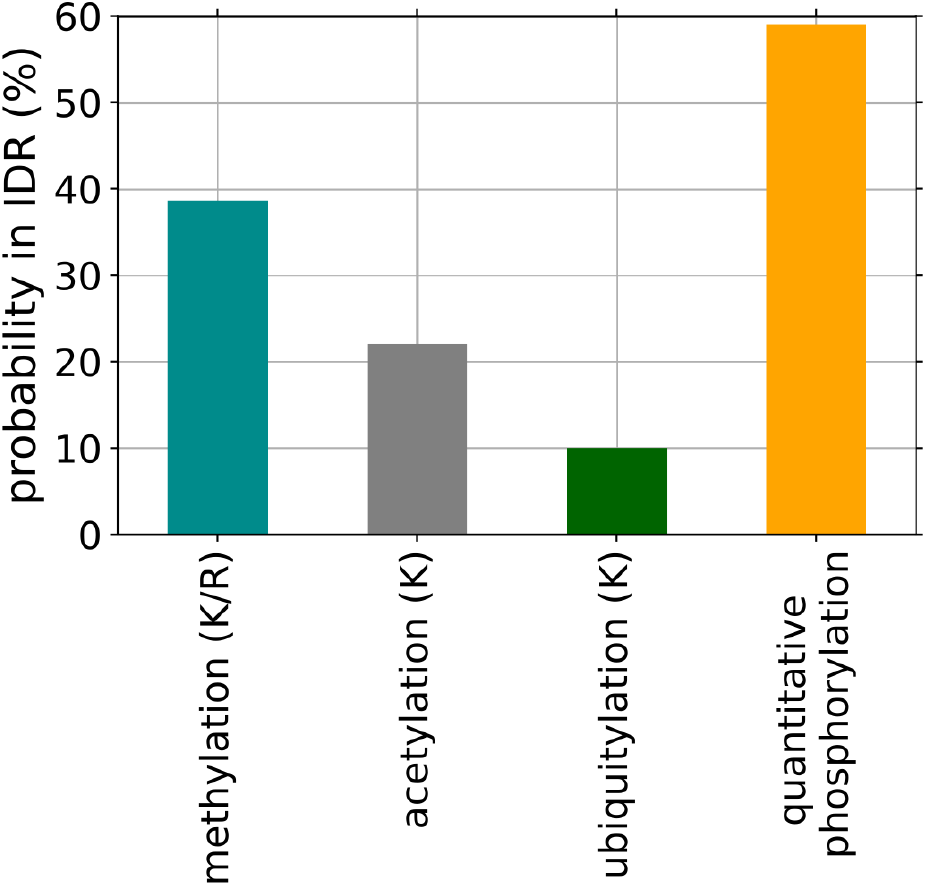
Probability of post-translational modifications occurred in IDR. Amino acid residues and its flanking amino acid sequences for methylation (15,283 sites), acetylation (20,875 sites) and ubiquitylation (96,796 sites) were extracted from PhosphositePlus, and were subjected to the analysis of IDR as described in Figure 2. As a comparison, the result of phosphosites obtained in this study is also plotted.

**Figure S4.**
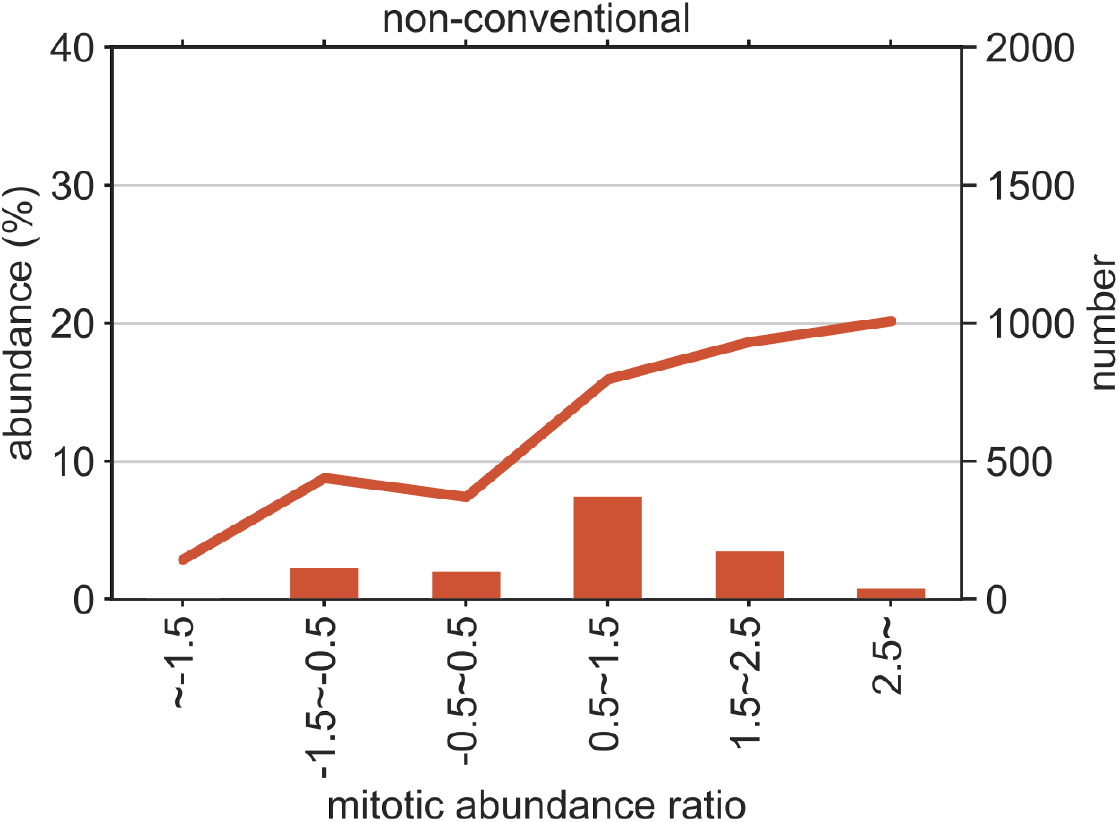
Phosphosites at non-conventional motif. The number of phosphosites at non-conventional motif (right axis) and its percentage (left axis) was plotted against mitotic abundance ratio.

**Table S1. Dataset from mass-spec analyses**

**Table S2. A list of proteins which carries NBC motifs and are known to be phosphorylated by AMPK.**

